# Discovering uncertainty: Bayesian constitutive artificial neural networks

**DOI:** 10.1101/2024.08.19.608595

**Authors:** Kevin Linka, Gerhard A Holzapfel, Ellen Kuhl

## Abstract

Understanding uncertainty is critical, especially when data are sparse and variations are large. Bayesian neural networks offer a powerful strategy to build predictable models from sparse data, and inherently quantify both, aleatoric uncertainties of the data and epistemic uncertainties of the model. Yet, classical Bayesian neural networks ignore the fundamental laws of physics, they are non-interpretable, and their parameters have no physical meaning. Here we integrate concepts of Bayesian learning and constitutive neural networks to discover interpretable models, parameters, and uncertainties that best explain soft matter systems. Instead of training an individual constitutive neural network and learning point values of the network weights, we train an ensemble of networks and learn probability distributions of the weights, along with their means, standard deviations, and credible intervals. We use variational Bayesian inference and adopt an efficient backpropagation-compatible algorithm that approximates the true probability distributions by simpler distributions and minimizes their divergence through variational learning. When trained on synthetic data, our Bayesian constitutive neural network successfully rediscovers the initial model, even in the presence of noise, and robustly discovers uncertainties, even from incomplete data. When trained on real data from healthy and aneurysmal human arteries, our network discovers a model with more stretch stiffening, more anisotropy, and more uncertainty for diseased than for healthy arteries. Our results demonstrate that Bayesian constitutive neural networks can successfully discriminate between healthy and diseased arteries, robustly discover interpretable models and parameters for both, and efficiently quantify uncertainties in model discovery. We anticipate our approach to generalize to other soft biomedical systems for which real-world data are rare and inter-personal variations are large. Ultimately, our calculated uncertainties will help enhance model robustness, promote personalized predictions, enable informed decision-making, and build confidence in automated model discovery and simulation.

Our source code, data, and examples are available at https://github.com/LivingMatterLab/CANN.

## 1. Motivation

When someone asks you *‘How confident are you in your neural network?’*, wouldn’t you want to respond *‘Very*.*’*? But what makes you believe that you are very confident? And how exactly do you quantify very? This is precisely what this manuscript is about.

Plain neural networks are prone to overfitting and non-interpretable [9]. They make overly confident decisions, generalize poorly, and are unsuitable for model discovery [40]. Here we address these limitations by using regularized variational Bayesian learning to *sparsify the weight vector* of a constitutive neural network [26] and *quantify the uncertainty* in the remaining non-zero weights [30]. The underlying idea is to represent the weights of a constitutive neural network by their probability distributions. Then, instead of training an *individual* constitutive neural network to learn point values of network weights [21], we train an *ensemble* of networks to learn probability distributions of the weights, with means, standard deviations, and credible intervals [25]. In other words, each network of the ensemble has its own network weights that we draw from shared probability distributions. We learn these probability distributions using Bayesian inference [2].

In practice, probability distributions of neural network weights can be quite complex, and, to complicate matters, the probabilities of individual weights can dependent on one another [3]. This makes, the *exact Bayesian inference* of the network weights a challenging if not infeasible task. Instead, we can numerically approximate Bayesian inference, and three classes of algorithms have emerged to do so: Markov Chain Monte Carlo methods that approximate posterior distribution of the network weights [30], dropout methods that probe discrete ensembles by setting subsets of weights to zero [8], and variational inference [10]. Here we use *variational Bayesian inference* [17] and adopt an efficient principled backpropagation-compatible algorithm [3] that approximates the probability distributions of our network weights with simpler distributions by minimizing their divergence through variational learning [12].

Bayesian neural networks are by no means new, in fact, they were first introduced in the early 1990s [25]. Within the science and engineering communities, they are enjoying increasingly popularity because they allow us to quantify both aleatoric and epistemic uncertainties [33]. *Aleatoric* or *data uncertainties* result from an inherent randomness or variability in the data, which is irreducible. *Epistemic* or *model uncertainties* result from a lack of knowledge or limitations in the model, which are potentially reducible, either by collecting more data or by refining the model [16]. We can quantify aleatoric uncertainties with both frequentist and Bayesian statistics, but only Bayesian statistics can quantify epistemic uncertainties [8]. It does so by treating the model parameters as probability distributions [15]. This introduces uncertainties in the model itself, which we could reduce by collecting more data.

In many science and engineering applications, clean data are rare and cumbersome to collect [1]. At the same time, we have a solid understanding of the underlying physics that our scientific community has built over many decades [14]. Naturally, this raises the question how to best build this knowledge into efficient learning machines [36]. Two different strategies have emerged to integrate physics-based knowledge into neural network models: *Physics Informed Neural Networks* or *PINNs* that incorporate physics into the loss function using additional terms [37], and *Constitutive Artificial Neural Networks* or *CANNs* that hardwire the underlying physics into the neural network design [19]. Our recent application of physics informed neural networks to real-world nonlinear dynamical systems has revealed the critical need to supplement these networks with uncertainty quantification, especially in situations where the underlying physics are not entirely known [20]. Here we build on this experience and explore the importance of uncertainty quantification in the context of constitutive artificial neural networks [21]. Our guiding question is: How can we discover the best model, parameters, and uncertainties for real-world nonlinear soft matter systems?

Towards this goal, in Section 2, we introduce the concept of Bayesian constitutive neural networks and derive their characteristic three-term loss function in the context of regularized variational Bayesian inference. In Section 3, we design a family of isotropic Bayesian constitutive neural networks and discover the best model, parameters, and uncertainties to explain *synthetic* soft matter data, perturbed by aleatoric noise. In Section 4, we generalize this concept towards transversely isotropic Bayesian constitutive neural networks and discover the best model, parameters, and uncertainties to explain *experimental* biaxial stress-stretch data of healthy and diseased human tissues. We compare our observations, discuss our results, and summarize our conclusions in Section 5.

## 2. Bayesian constitutive neural networks

In the following, we briefly summarize the concept of Bayesian inference [2], discuss its computational realization using variational Bayesian inference [10], and derive the loss function for Bayesian constitutive neural networks towards automated model discovery [21].

### Bayesian inference

Bayesian inference is a statistical method that updates the probability for a hypothesis as more information becomes available using Bayes’ theorem. Bayes’ theorem states that the posterior probability is equal to the likelihood times the prior probability, divided by the marginal likelihood or evidence,

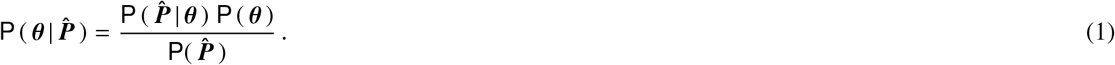

Here 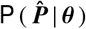 is the *likelihood function*, in our case the conditional probability of the measured stresses 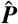 for given fixed parameters ***θ***; `P` (***θ***) is the *prior probability distribution* of the model parameters 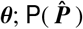 is the *marginal likelihood* or *evidence*; and 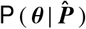 is the *posterior probability distribution*, the conditional probability of the parameters ***θ*** for given measured stresses 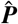. We can use Bayes’ theorem to infer the posterior probability distribution, and with it the model, parameters, and uncertainty that best explain given data, in our case, labeled stress-stretch pairs.

### Likelihood

The *likelihood* function 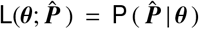 quantifies the chance that some calculated parameters ***θ*** explain the measured stress 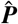. It measures the goodness of fit between the observed stress-stretch data 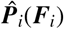 and the model output ***P***(***θ***, ***F***_*i*_), the discovered stress-stretch model of the neural network for the learnt parameters ***θ***, at a fixed deformation gradient ***F***_*i*_. The overall likelihood 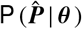 is the product of *i* = 1, …, n individual likelihood functions 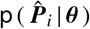, one for each deformation state ***F***_*i*_. A common choice for 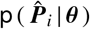 is the normal distribution for which the measurements are centered around the mean *µ*_*i*_ at the hidden real values 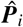, with standard deviations *σ*_*i*_ that account for measurement errors, i.e.,

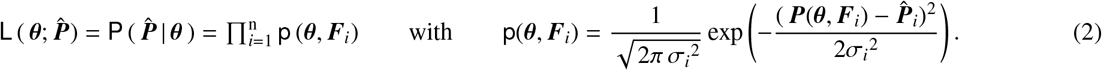

A well-known disadvantage of the likelihood function (2) is that it involves the product of *i* = 1, …, n probabilities, which can become extremely small, especially when dealing with large datasets. Computationally, too small likelihoods can result in numerical instabilities as their product becomes too close to zero. Many machine learning tools avoid the product of probabilities in the likelihood function (2) by using the natural logarithm of the likelihood, the *log-likelihood* function 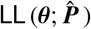 instead,

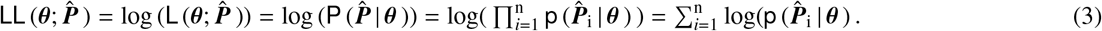

The log-transformation translates the product of probabilities into a sum of log-probabilities and helps maintain computational precision and numerical stability. Computationally, it often proves convenient to convert the log-likelihood function (3) into the *negative log-likelihood* function 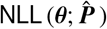,

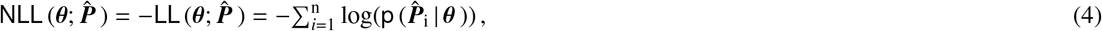

and minimize the negative log-likelihood `NLL` rather than maximize the log-likelihood `LL`.

### Maximum likelihood estimate

The parameter values that maximize the log-likelihood function 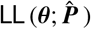 from equation (3) are the *maximum likelihood estimate* of the true parameter values ***θ***,

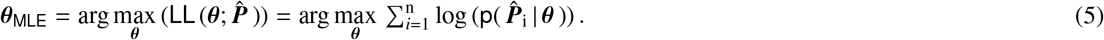

Intuitively, the maximum likelihood estimate ***θ***_MLE_ defines the set of parameters values that make the observed data, in our case the measured stresses 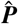, most probable. Maximizing the log-likelihood 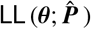 from equation (3) is equivalent to minimizing the negative log-likelihood 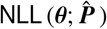 from equation (4). This implies that the parameter vector that *maximizes the log-likelihood **θ***_MLE_ is also the parameter vector that *minimizes the negative-log-likelihood*,

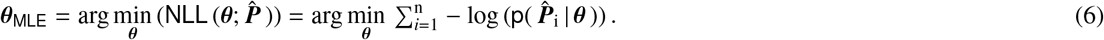

Many machine learning tools have built-in algorithms for minimization, and favor the definition (6) over (5).

### Priors

The *prior probability distribution* `P` (***θ***) represents our initial belief about the distribution of the parameters before we have observed any data. In our case, the model parameters are the weights of the neural network, ***θ*** = { ***w, w***^∗^ }. We can distinguish two types of network weights, the *external* weights ***w*** out of the last hidden layer and the *internal* weights ***w***^∗^ between the hidden layers. For each parameter we would like to infer, we have to select a prior probability distribution 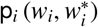 and the total prior probability density `P` (***θ***) becomes the product of all *i* = 1, …, n individual distributions,

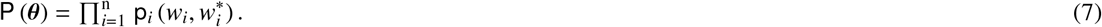

To increase the robustness of our model discovery, we only apply prior probability distributions to the external weights ***w*** = { *w*_*i*_ }, while keeping the internal weights 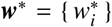 deterministic. The choice of the individual distributions `p`_*i*_ is based on our prior domain knowledge or simply on mathematical convenience. For example, we could choose Gaussian distributions with probability densities

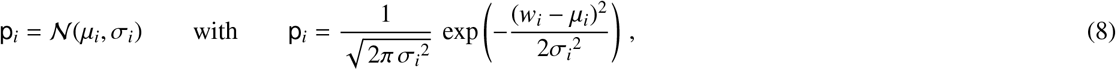

with fixed means *µ*_*i*_ and standard deviations *σ*_*i*_, or uniform distributions `p`_*i*_ = 1 / (*w*_*i*,max_ − *w*_*i*,min_) with fixed upper and lower bounds *w*_*i*,max_ and *w*_*i*,min_. In Bayesian statistics, we use the prior `P`(***θ***) to update the beliefs about the parameters ***θ*** after observing the data 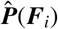 using Bayes’ theorem (1). Importantly, while the priors can take various forms, such as normal or uniform, they are *fixed* distributions that represent our *initial* beliefs about the parameters.

### Marginal likelihood

The *marginal likelihood* function 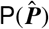 is also often referred to as evidence. It represents a likelihood function in which the parameter variables are marginalized by integrating over the entire parameter domain,

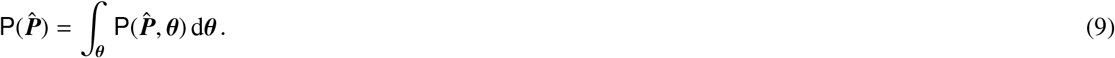

Marginal likelihoods are generally difficult if not impossible to compute. If needed, we could integrate equation (9) using numerical integration schemes such as Gaussian integration or Monte Carlo methods. Fortunately, for most practical purposes, we do not need to know the precise value of the marginal likelihood 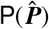 Instead of evaluating *absolute* probabilities using Bayes’ theorem (1), we typically focus on *relative* probabilities using the ratio of two posterior probabilities, 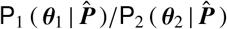 Since the marginal likelihood is a normalizing constant, it cancels out when computing relative probabilities. For model comparison and parameter estimation tasks, we often reduce Bayes’ theorem (1) to the fact that the posterior distribution 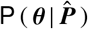 is proportional to the product of the likelihood and the prior, i.e.,

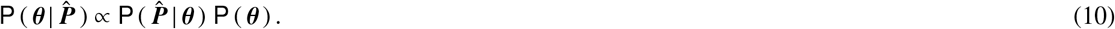

For most practical purposes, knowing this proportionality (10) is sufficient and we do not need to know the exact value of the marginal likelihood 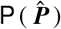

### Posterior

The *posterior probability distribution* 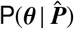 is the conditional probability of the parameters ***θ*** for the given data 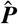, the experimentally measured stress. We can calculate the posterior probability from the likelihood 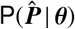 and our prior knowledge encoded through the prior probability distribution `P`(***θ***), using Bayes’ theorem (1),

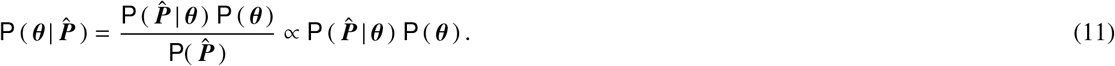

The parameters ***θ*** that maximize the posterior probability distribution 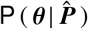 are called the *maximum a posteriori estimate **θ***_MAP_ of the true parameter values ***θ***, and their values are independent of the marginal likelihood 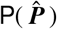,

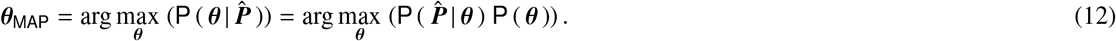

A common strategy to estimate the posterior distributions (11) and the maximum a posteriori estimate (12) is multi-chain full-batch Hamiltonian Monte Carlo, a highly efficient and well-studied Markov Chain Monte Carlo method. Theoretically, Hamiltonian Monte Carlo is guaranteed to produce samples from the true posterior asymptotically. In practise, applying Hamiltonian Monte Carlo to state-of-the-art neural networks is extremely challenging due to its high computational cost: It can take tens of thousands of training epochs to produce a single sample from the posterior.

### Variational Bayesian inference

Variational Bayesian inference has become a popular method to speed-up computation when estimating complex probability distributions in the context of Bayesian statistics. Its underlying idea is to *approximate complex posterior distributions* with a much simpler distribution, the *variational distribution*. Here we approximate the posterior probability distribution 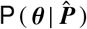 of the unknown parameter vector ***θ*** = {***w, w***^∗^} by a less flexible family of distributions, the variational distribution `Q` (***θ***; ***W***) of the parameter vector ***W*** = {***w, w***^∗^, ***w***_*µ*_, ***w***_*σ*_}. This new parameter vector ***W*** consists of two types of parameters, the *network weights* ***w*** and ***w***^∗^, and the *variational parameters* ***w***_*µ*_ and ***w***_*σ*_. We follow the standard approach [3] and assume that the variational approximation `Q` (***θ***; ***W***) adopts a parameterized Gaussian distribution,

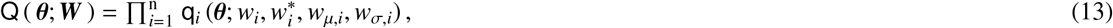

as product of *i* = 1, …, n one-dimensional Gaussian distributions `q`_*i*_. Each individual Gaussian distribution is the product of the normal distribution, 𝒩(*w*_*µ,i*_, *w*_*σ,i*_), in terms of the mean and standard deviation *w*_*µ,i*_ and *w*_*σ,i*_, and the neural network activation function 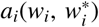 in terms of the external and internal weights *w*_*i*_ and 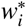,

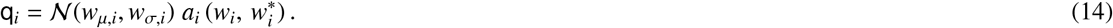

Importantly, instead of using *fixed* means and standard deviations *µ*_*i*_ and *σ*_*i*_ as in equation (8), variational inferences treats the means and standard deviations *w*_*µ,i*_ and *w*_*σ,i*_ as *trainable* variational parameters. Yet, here, instead of using the standard deviation *w*_*σ,i*_ directly, we subject it to the exponential linear unit function `elu`(*w*_*σ,i*_), with `elu`(○) = (○) for non-negative arguments (○) ≥ 0, and `elu`(○) = *α*(exp((○) − 1) for negative arguments (○) < 0. Using this modified distribution, 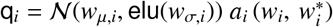 instead of equation (14) manages negative values by pushing them closer to zero and speeds up learning by bringing the normal gradient closer to the unit natural gradient [4]. Now, the objective is to find the optimal parameters ***W***, such that the variational approximation `Q`(***θ***; ***W***) is as close to the true posterior distribution 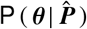 as possible. We measure the divergence between `Q`(***θ***; ***W***) and 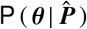 using the *Kullbach-Leibler divergence*,

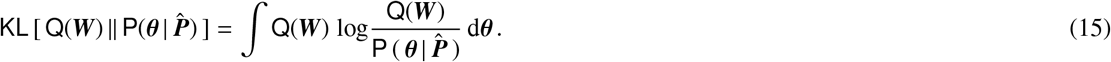

With the definition of the conditional probability, 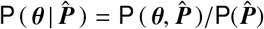, and the calculus rules of the logarithm, log(*A* · *B*) = log(*A*) + log(*B*) and log(*A*/*B*) = − log(*B*/*A*), this definition becomes

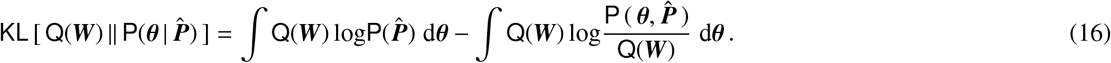

Since the probability of the data 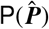 is independent of the parameters ***θ***, the first integral simplifies to 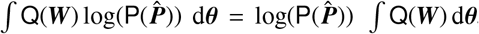, where the integral for all probability densities `Q`(***W***) is identical to one, ∫ `Q`(***W***) d***θ*** = 1, and the Kullbach-Leibler divergence (15) is identical to the following expression,

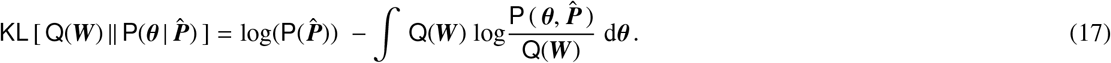

### Minimum Kullbach-Leibler divergence

The objective of the variational inference is to find the parameters ***W*** that minimize the Kullbach-Leibler divergence (17). Since the probability of the data 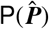 is independent of the parameters ***W***, we only need to minimize the second term,

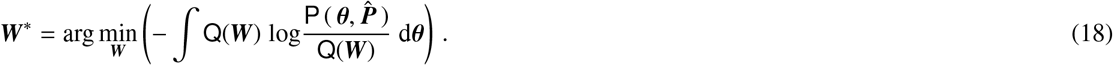

We now eliminate the unknown posterior probability using the definition of the conditional probability, 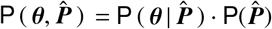, and apply the calculus rules of the logarithm,

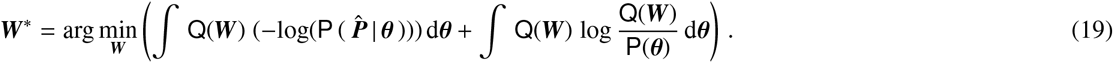

The first term is the expected negative log-likelihood, 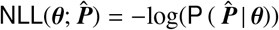, given the parameters ***θ*** distributed according to the variational distribution `Q`, and the second term is the Kullbach-Leibler divergence, 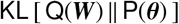,

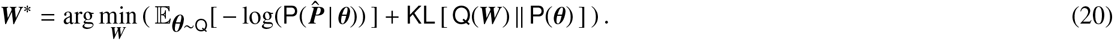

We now formulate the loss function for our Bayesian constitutive neural network motivated by equation (20).

### Loss function

We train our Bayesian constitutive neural network by minimizing a three-term loss function *L* to learn the network weights ***w*** and ***w***^∗^ and the variational parameters ***w***_*µ*_ and ***w***_*σ*_ [3], i.e.,

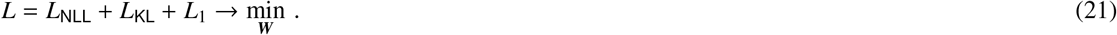

The first term represents the expected negative log-likelihood `NNL` according to equation (4) given the parameters ***θ*** with the variational distribution `Q`,

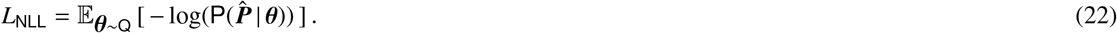

The second term is the Kullback-Leibler divergence between the variational approximation `Q`(***θ***; ***W***) that approximates the parameters ***θ*** using ***W*** and the prior `P`(***θ***). By minimizing this second term, variational learning efficiently approximates the prior distribution `P`(***θ***) when the exact distribution is unknown,

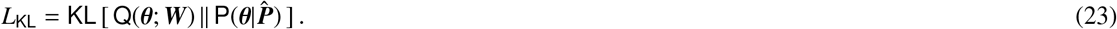

The third term is the *L*_1_ regularization that sparsifies the model by penalizing the weighted *L*_1_ norm of the external weights ***w*** [41], where the penalty parameter *α* determines the number of active terms [26],

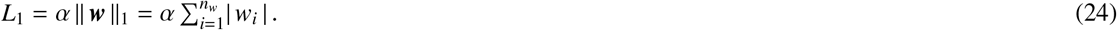

In summary, the loss function (21) is a sum of a data-dependent part associated with the likelihood cost, a prior dependent part associated with the complexity cost, and a regularization-dependent part associated with the sparsity cost [3]. We implement our Bayesian constitutive neural networks in Tensorflow-Probability and minimize the loss function using the ADAM optimizer, a robust adaptive algorithm for gradient-based first-order optimization. Figure 1 illustrates the convergence of the loss function during a representative training run. Both the Kulbach-Leibler divergence *L*_KL_ and the total loss function *L* = *L*_NLL_ + *L*_KL_ + *L*_1_ converge robustly, here for an example of Section 3 within 500 epochs, as the network learns the network weights ***w*** and ***w***^∗^ and the variational parameters ***w***_*µ*_ and ***w***_*σ*_.

**Figure 1:**
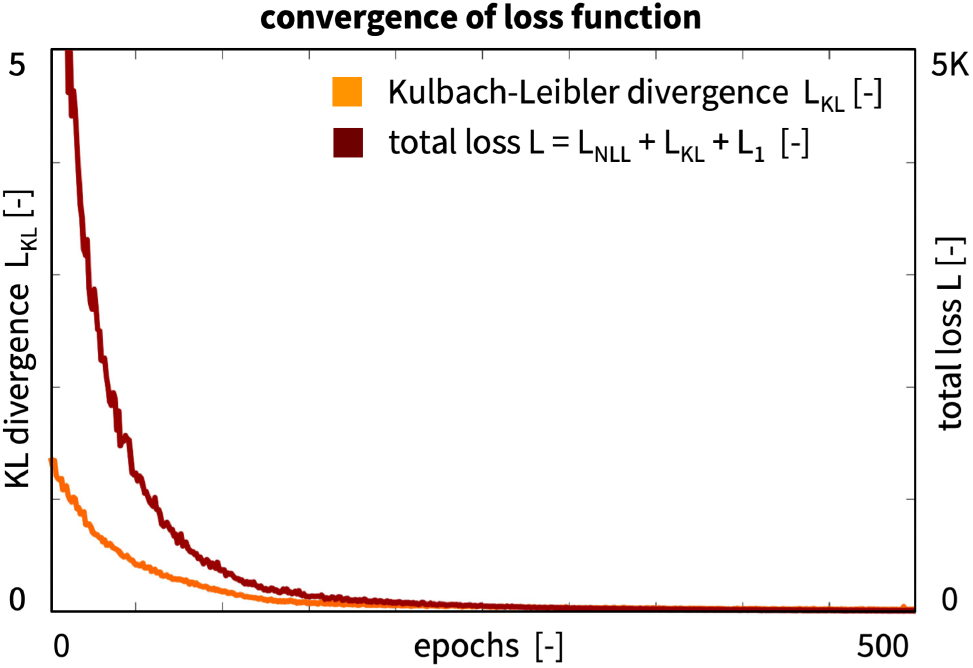
Convergence of loss function. During a representative training run, the Kulbach-Leibler divergence *L*_KL_ and the total loss function *L* = *L*_NLL_ + *L*_KL_ + *L*_1_ converge robustly as the network learns the network weights ***w*** and ***w***^∗^ and the variational parameters ***w***_*µ*_ and ***w***_*σ*_.

From the learnt variational approximation `Q` (***θ***; ***W***) of the true posterior distribution 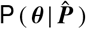, we sample *i* = 1, …, *M* model parameters 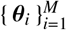 and derive the stresses 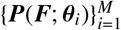 for each sample. We report the means and standard deviations of the stresses, where the former represent the model prediction and the latter quantify the model uncertainty.

In the following, we demonstrate how to design Bayesian constitutive neural networks for *isotropic* and *transversely isotropic* incompressible hyperelastic materials. For the isotropic case, we discover the best model, parameters, and posterior distributions of the stresses to explain *synthetic* stress-stretch data of a Mooney-Rivlin material in uniaxial tension, uniaxial compression, equibiaxial tension, and pure shear, perturbed by aleatoric noise. For the transversely isotropic case, we discover the best model, parameters, and posterior distributions of the stresses to explain *experimental* stress-stretch data of healthy and diseased abdominal aortic tissue from the medial layer and from both media and adventitia layers combined when tested in equibiaxial tension.

## 3. Isotropic Bayesian constitutive neural networks

The *isotropic* Bayesian constitutive neural network takes the deformation gradient ***F***, the gradient of the deformation map ***φ*** with respect to the coordinates of the undeformed sample ***X***, as input,

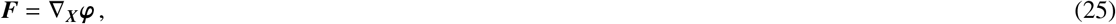

and calculates the first and second invariants,

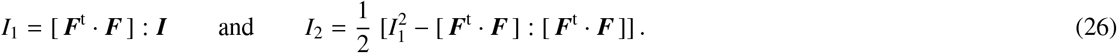

For *perfectly incompressible* materials, the third invariant remains constant and identical to one, *I*_3_ = *J*^2^ = 1. The network discovers *hyperelastic* material models that satisfy the second law of thermodynamics, which implies that the Piola stress ***P*** is the derivative of the free energy *ψ*(***F***) with respect to the deformation gradient ***F*** modified by a pressure term, −*p* ***F***^−t^,

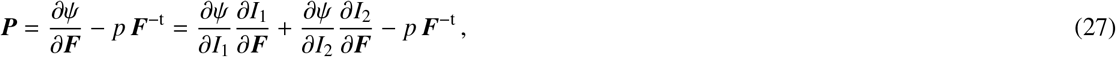

where the hydrostatic pressure, 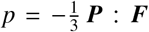, acts as a Lagrange multiplier that we determine from the boundary conditions. We discover the free-energy function *ψ* using a Bayesian constitutive neural network that takes the deformation gradient ***F*** as input and approximates the free-energy function *ψ*(***F***) as the sum of eight probability-weighted terms. Figure 2 illustrates our neural network with two hidden layers and eight nodes [21]. The first layer generates powers (○) and (○)^2^ of the network input, the two invariants *I*_1_ and *I*_2_. The second layer applies the identity, (○) and the exponential function (exp(○)) to these powers. The network output is the sum of these eight terms, weighted by their probability densities 𝒩(*w*_*µ*_, *w*_*σ*_). Importantly, since we focus on model discovery, we only introduce probabilistic external weights, while keeping all internal weights deterministic. This naturally limits the number of additional parameters, and makes the network more robust by design. The free-energy function of this networks takes the following explicit form,

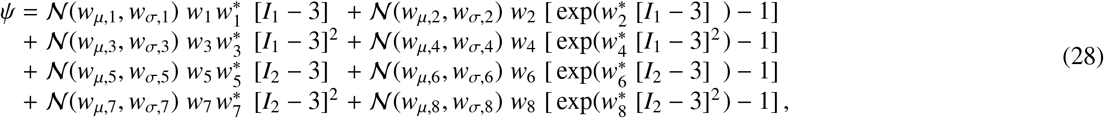

corrected by the pressure term, *ψ* = *ψ* − *p* [*J* − 1]. To complete the definition of the Piola stress in equation (27), we take its derivatives with respect to the two invariants,

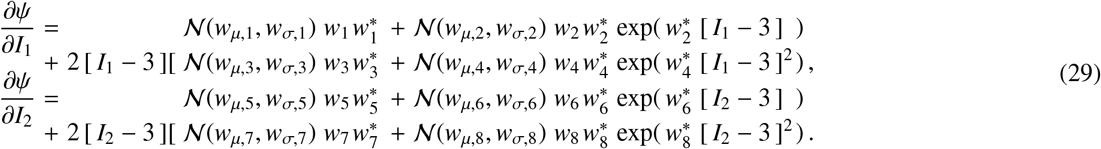

**Figure 2:**
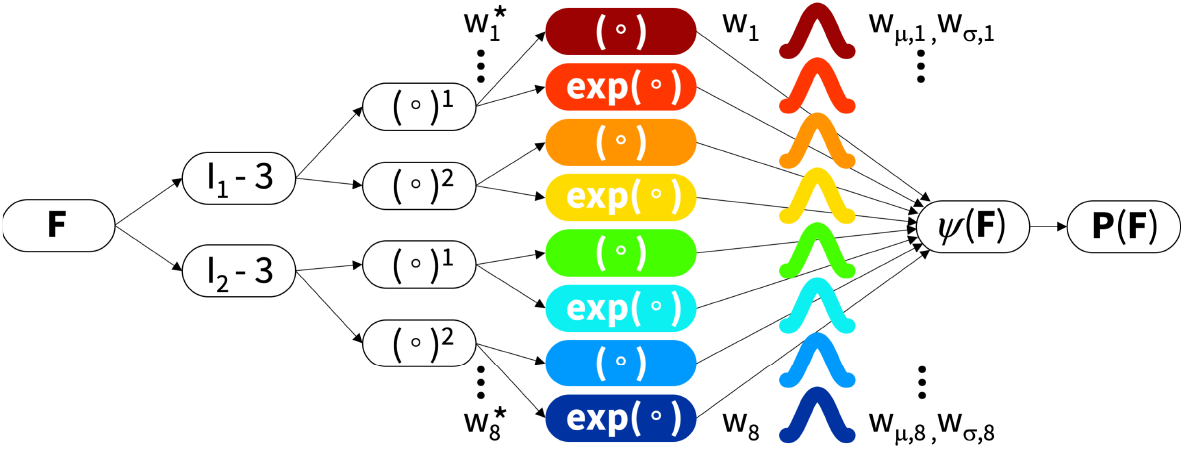
Isotropic Bayesian constitutive neural network. The network has two hidden layers to discover the free-energy function *ψ*(*I*_1_, *I*_2_) as a function of the invariants of the deformation gradient ***F*** using eight terms. The first layer generates powers (○) and (○)^2^ of the network input, the second layer applies the identity (○) and exponential function (exp(○)) to these powers, and the network output is the sum of these eight terms, weighted by their probability densities 𝒩(***w***_*µ*_, ***w***_*σ*_). During training, the network learns the network weights ***w*** and ***w***^∗^ and the variational parameters ***w***_*µ*_ and ***w***_*σ*_.

The network has two sets of network weights, ***w*** and ***w***^∗^, associated the eight terms of the free-energy function, and two sets of variational parameters, ***w***_*µ*_ and ***w***_*σ*_, associated with the means and standard deviations of these eight terms. We learn these 32 weights by minimizing the three-term loss function *L* = *L*_NLL_ + *L*_KL_ + *L*_1_ in equation (21).We consider data from homogeneous uniaxial compression, uniaxial tension, equibiaxial tension, and pure shear test.

For the cases of *uniaxial compression*, and *uniaxial tension*, with a stretch *λ* in the 1-direction, such that *I*_1_ = *λ*^2^ +2*λ*^−1^ and *I*_2_ = 2*λ* + *λ*^−2^ and **F** = diag { *λ, λ*^−1/2^, *λ*^−1/2^ } and **P** = diag { *P*_11_, 0, 0 }, the stress-stretch relation for isotropic materials [22] is

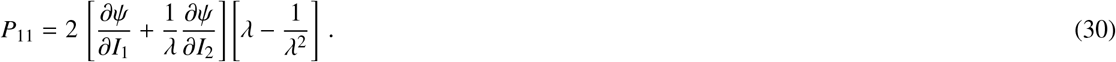

For the case of *equibiaxial tension*, with a stretch *λ* in the 1- and 2-directions, such that *I*_1_ = 2*λ*^2^+*λ*^−4^ and *I*_2_ = *λ*^4^+2*λ*^−2^ and **F** = diag { *λ, λ, λ*^−2^ } and **P** = diag { *P*_11_, *P*_22_, 0 }, the stress-stretch relation for isotropic materials [21] is

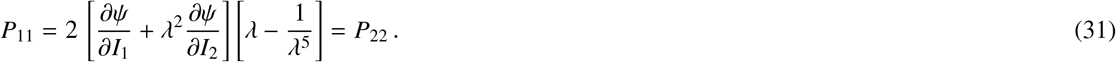

For the case of *pure shear* of a long rectangular specimen stretched with *λ* along its short axis in the 1-direction, and no deformation along it long axis in the 2-direction, such that *I*_1_ = *I*_2_ = *λ*^2^ + 1 + *λ*^−2^ and **F** = diag { *λ*, 1, *λ*^−1^ } and **P** = diag { *P*_11_, *P*_22_, 0 }, the stress-stretch relations for isotropic materials [21] are

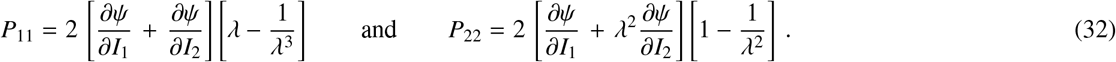

We explore the performance of the isotropic Bayesian neural network from Figure 2 on the basis of synthetic stress-stretch data of a Mooney-Rivlin material [29, 38] perturbed by aleatoric noise. The strain energy function of the Mooney-Rivlin model is 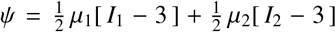, where *µ*_1_ and *µ*_2_ denote the two model parameters. Following equations (30), (31) and (32), we obtain the explicit stress-stretch relationships for uniaxial compression and tension, *P*_11_ = [ *µ*_1_ + *λ*^−1^*µ*_2_ ] [ *λ* − *λ*^−2^ ], for equibiaxial tension, *P*_11_ = [ *µ*_1_ + *λ*^2^*µ*_2_ ] [ *λ* − *λ*^−5^ ], and for pure shear, *P*_11_ = [ *µ*_1_ + *µ*_2_ ] [ *λ* − *λ*^−3^ ]. To generate the synthetic data, we apply an additive Gaussian aleatoric noise to the stress, 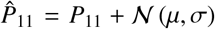. We select a zero mean, *µ* = 0, and a standard deviation that scales linearly with the absolute stress, *σ* = *σ*_0_ | *P*_11_ |, and sample synthetic stress-stretch data pairs 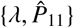 for all load cases. We then apply Bayesian model discovery: We train the network by minimizing the loss function *L*, draw parameter samples 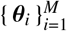 from the learnt variational approximation `Q` (***θ***; ***W***) of the true posterior distribution 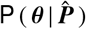, derive stresses 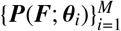 for each sample from equations (30), (31), (32), and report the means and standard deviations of the stresses. To explore the predictive potential of the Bayesian network and evaluating model robustness, we compare two cases, training on *all* data and training on *incomplete* data while testing on the remaining data.

### Bayesian networks successfully rediscover the initial model, even in the presence of noise

Figure 3 shows the discovered model for the synthetic data of the Mooney-Rivlin model, 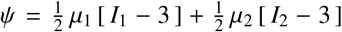, with true parameters *µ*_1_ = 1 kPa and *µ*_2_ = 1 kPa and an aleatoric scaling coefficient *σ*_0_ = 0.15, and an *L*_1_ regularization parameter *α* = 0.001, trained on *all* data in the stretch ranges of *λ* = [ 0.8, …, 1.0 ] for uniaxial compression, and *λ* = [ 1.0, …, 1.3 ] for uniaxial tension, equibiaxial tension, and pure shear. Figure 4 highlights the discovered isotropic model parameters along with their posterior distributions. Both figures confirm that our Bayesian constitutive neural network successfully identifies the mean stress response and the uncertainty in the data. First and foremost, out of 2^8^ = 256 possible combinations of terms, the network robustly rediscovers the Mooney Rivlin model, 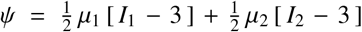 with mean network weights of *w*_1_ = 0.782 kPa and *w*_5_ = 0.525 kPa, which translate into Mooney Rivlin parameters of *µ*_1_ = 2 *w*_1_ = 1.564 kPa and *µ*_2_ = 2 *w*_5_ = 1.050 kPa, two meaningful parameters with physical units and a clear physical interpretation [29, 38]. In addition, the network also discovers the posterior distributions of the weights along with their standard deviations, *w*_1_ = 0.782 ± 0.02 kPa and *w*_5_ = 0.525 ± 0.01 kPa. Most importantly, the network autonomously trains all inactivate model terms to zero [26], *w*_2_ = *w*_3_ = *w*_4_ = *w*_6_ = *w*_7_ = *w*_8_ = 0 kPa, which is not straightforward in conventional Bayesian neural networks [12]. Taken together, our Bayesian constitutive neural networks robustly and repeatably discover models, parameters, and uncertainty from synthetic data, even in the presence of noise.

**Figure 3:**
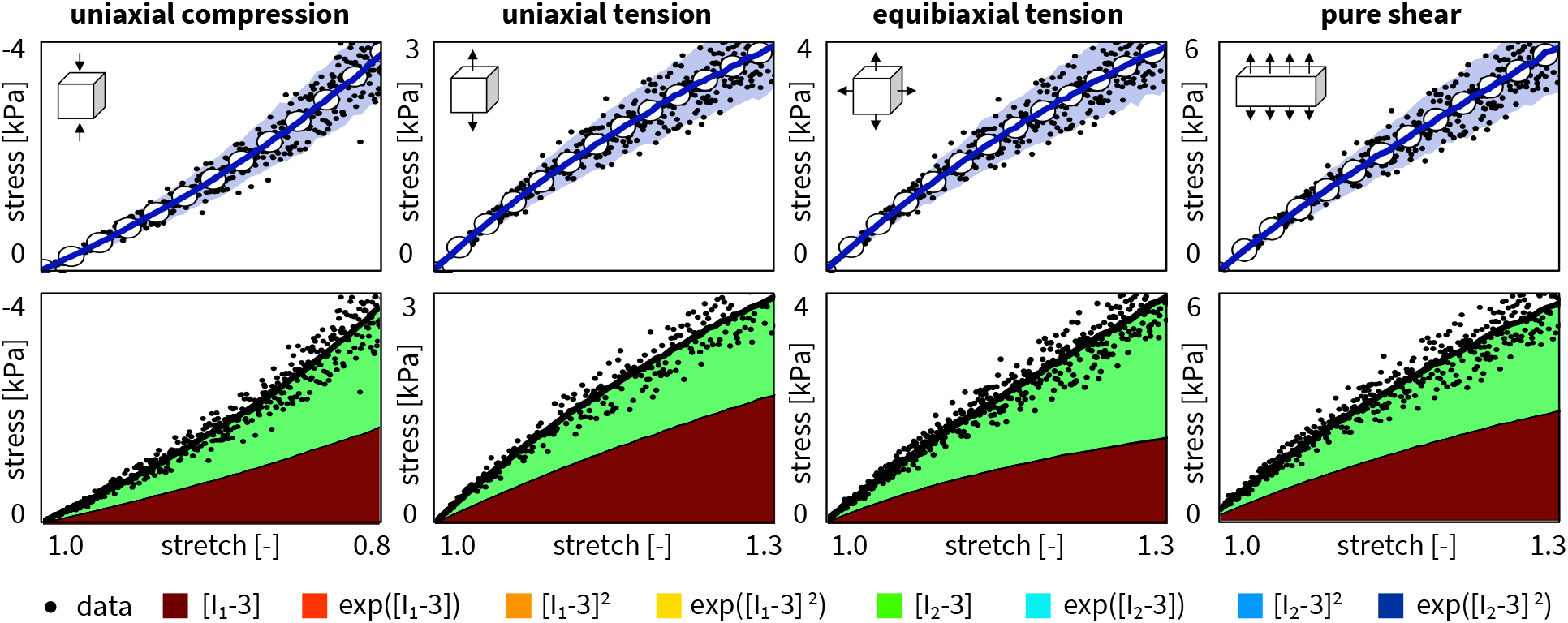
Synthetic data and discovered isotropic model for training on all data. Nominal stresses P as functions of the stretches *λ* for the isotropic, perfectly incompressible Bayesian constitutive neural network with two hidden layers, eight nodes, and eight prior distributions from Figure 2. We used all compressive data up to *λ* = 0.8, and all tensile, biaxial, and shear data up to *λ* = 1.3 for training. Dots illustrate the synthetic uniaxial compression, uniaxial tension, equibiaxial tension, and pure shear data of the Mooney Rivlin model with Gaussian noise; solid blue curves and blue-shaded areas indicate the mean predicted stresses ± standard deviations; color-coded areas highlight the eight possible contributions to the discovered stress function.

**Figure 4:**
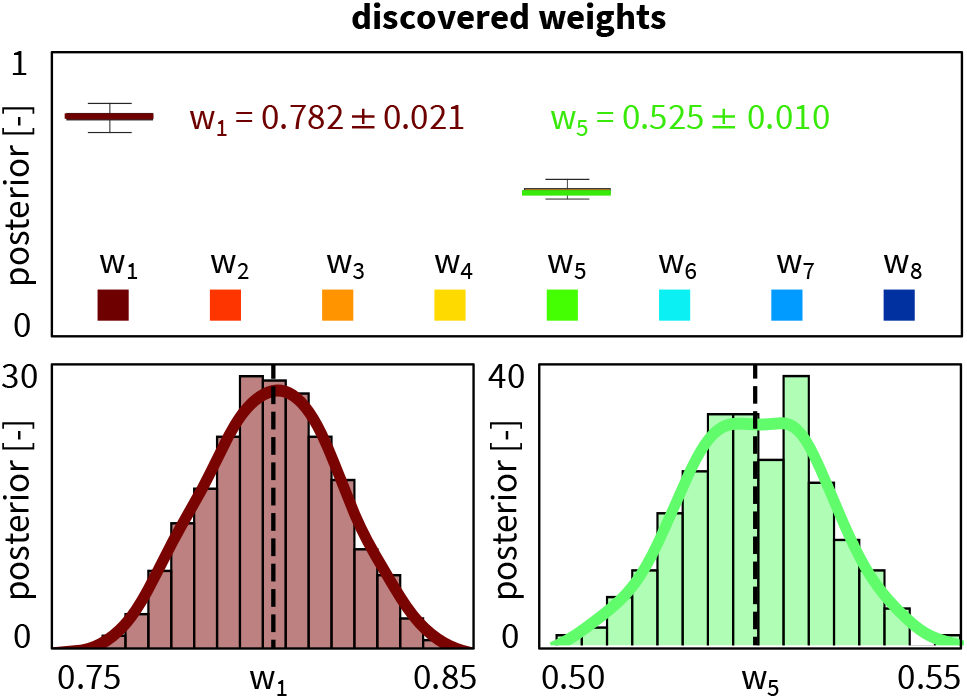
Posterior distributions of discovered isotropic model parameters for training on all data. When using all data for training, the isotropic, perfectly incompressible Bayesian constitutive neural network robustly rediscovers the Gaussian noise perturbed Mooney Rivlin model, *ψ* = *w*_1_ [ *I*_1_ − 3 ] + *w*_5_ [ *I*_2_ − 3 ], with discovered network weights of *w*_1_ = 0.782 ± 0.02 and *w*_5_ = 0.525 ± 0.01, while all other network weights correctly train to zero.

### Bayesian networks robustly discover uncertainties in non-training regimes

Figure 5 shows the discovered model for the synthetic data of the Mooney-Rivlin model, 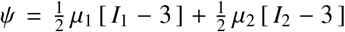, with true parameters *µ*_1_ = 1 kPa and *µ*_2_ = 1 kPa, an aleatoric scaling coefficient *σ*_0_ = 0.15, and an *L*_1_ regularization parameter *α* = 0.001, trained on *incomplete* data in the stretch ranges of *λ* = [ 0.8, …, 1.0 ] for uniaxial compression, and *λ* = [ 1.0, …, 1.15 ] for uniaxial tension, equibiaxial tension, and pure shear, and tested in the unseen stretch rage of *λ* = [ 1.15, …, 1.3 ]. Figure 6 highlights the discovered isotropic model parameters along with their posterior distributions. Similar to the previous example, our Bayesian constitutive neural network successfully discovers the mean stress response and the uncertainty in the data. However, the direct comparison of the uncertainties in the stress predictions in the blue-shaded areas in Figures 3 and 5 confirms our intuition that training on *all* data in Figure 3 results in smaller uncertainties than training on *incomplete* data in Figure 5. Interestingly, the extrapolation into the unseen regime of 1.15 < *λ* ≤ 1.3 is better in uniaxial tension and pure shear than in equibiaxial tension, where the uncertainties increase dramatically as the stretch increases. Nonetheless, the network is still able to discover the initial Mooney Rivlin model [29, 38], 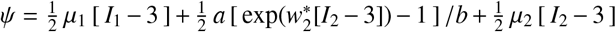, but now with an additional exponential linear first invariant Demiray-type term [5], 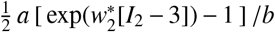 The discovered mean network weights of *w*_1_ = 0.824 kPa, *w*_2_ = 0.054 kPa and *w*_5_ = 0.468 kPa translate into Mooney Rivlin stiffnesses of *µ*_1_ = 2 *w*_1_ = 1.648 kPa and *µ*_2_ = 2 *w*_5_ = 0.936 kPa, a stiffness like parameter of *a* = 2 *w*_2_ *w*_2_ = 1.42 kPa, and a nonlinearity parameter of 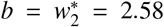, four meaningful parameters with physical units and a clear physical interpretation. In addition, the network also discovers the posterior distributions of the weights along with their standard deviations, *w*_1_ = 0.824 ± 0.010 kPa, *w*_2_ = 0.054 ± 0.010 kPa, and *w*_5_ = 0.468 ± 0.003 kPa. We observe that the non-Mooney Rivlin weight *w*_2_ = 0.054 is an order of magnitude smaller than the two Mooney Rivlin weights *w*_1_ = 0.824 and *w*_5_ = 0.468. We conclude that the influence of the additional exponential linear first invariant Demiray-type term [5], 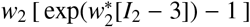 is relatively small, and that the discovered model remains dominated by the two Mooney Rivlin terms [29, 38], *w*_1_ [ *I*_1_ − 3 ] and *w*_5_ [ *I*_2_ − 3 ]. The small narrow red band of the exponential linear first invariant contribution to the overall stress in Figure 5 confirms this observation. Increasing the penalty parameter *α* beyond *α* = 0.001 would reduce the number of active model terms. Naturally, the smallest weight, *w*_2_ = 0.054, would be dropped first, and we would recover the original Mooney Rivlin model. Importantly, even for incomplete training data, the network autonomously identifies the true non-zero model terms *w*_1_ and *w*_5_, while training all other weights, except the additional small *w*_2_ term, to zero, *w*_3_ = *w*_4_ = *w*_6_ = *w*_7_ = *w*_8_ = 0 kPa. We have confirmed, although not explicitly shown here, that we can explicitly modulate the sparsification of the parameter vector ***w*** through the penalty parameter *α* [11] to fine-tune the trade-off between model accuracy and model interpretability [7]. Especially in the context of interpretability, our specific network architecture, with standard external and internal network weights ***w*** and ***w***^∗^ and variational parameters ***w***_*µ*_ and ***w***_*σ*_ only *after* the final hidden layer, turns out to be critical to selectively modulate sparsification. While we did experiment with alternative network architectures in which the variational parameters *replace* the external network weights ***w***, or all external and internal weights ***w*** and ***w***^∗^ [30], we believe that our current network architecture in Figure 3 provides the most robust parameter sparsification and the most interpretable model discovery. Taken together, our Bayesian constitutive neural network robustly and repeatably discovers predictive models, parameters, and uncertainty from synthetic data, even in the presence of noise and incomplete training data.

**Figure 5:**
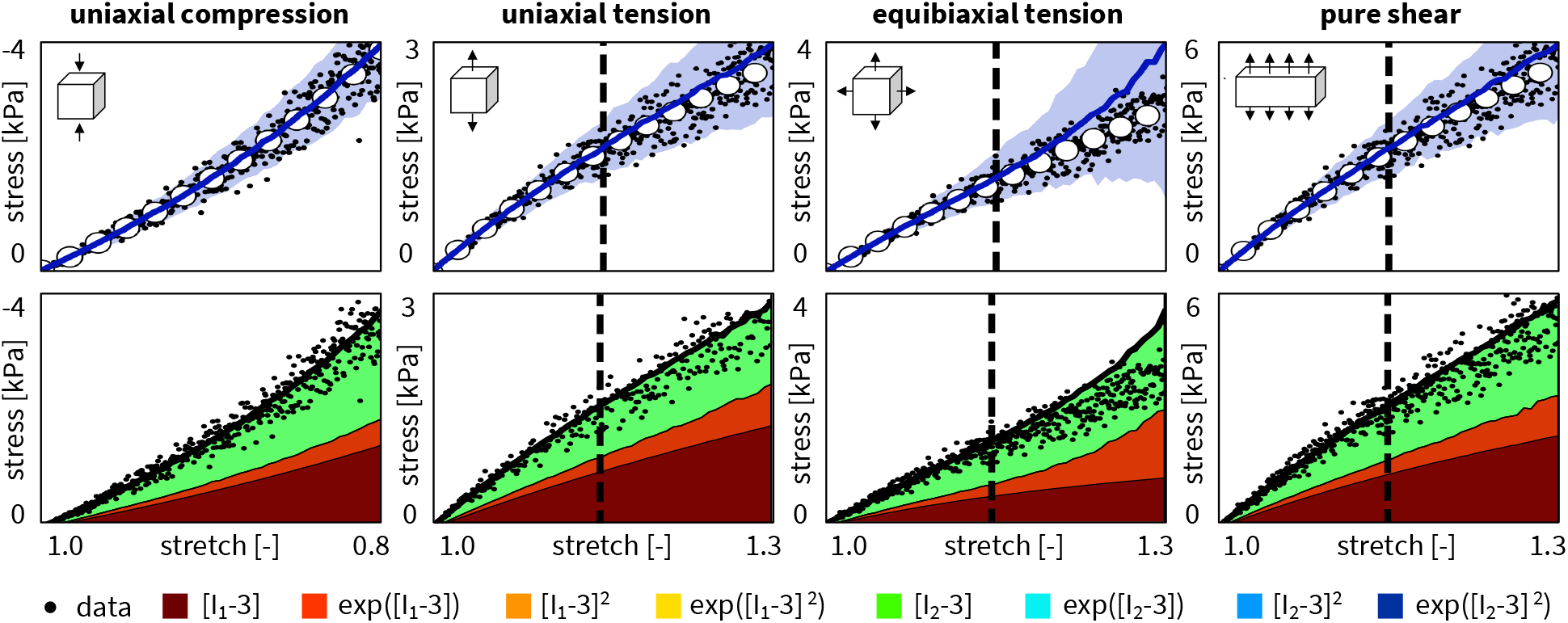
Synthetic data and discovered isotropic model for training on incomplete data. Nominal stresses *P* as functions of the stretches *λ* for the isotropic, perfectly incompressible Bayesian constitutive neural network with two hidden layers, eight nodes, and eight prior distributions from Figure 2. We used all compressive data up to *λ* = 0.8 for training, and tensile, biaxial, and shear data up to *λ* = 1.15 for training, and up to *λ* = 1.3 for testing. Dots illustrate the synthetic uniaxial compression, uniaxial tension, equibiaxial tension, and pure shear data of the Mooney Rivlin model with Gaussian noise; solid blue curves and blue-shaded areas indicate the mean predicted stresses ± standard deviations; color-coded areas highlight the eight possible contributions to the discovered stress function.

**Figure 6:**
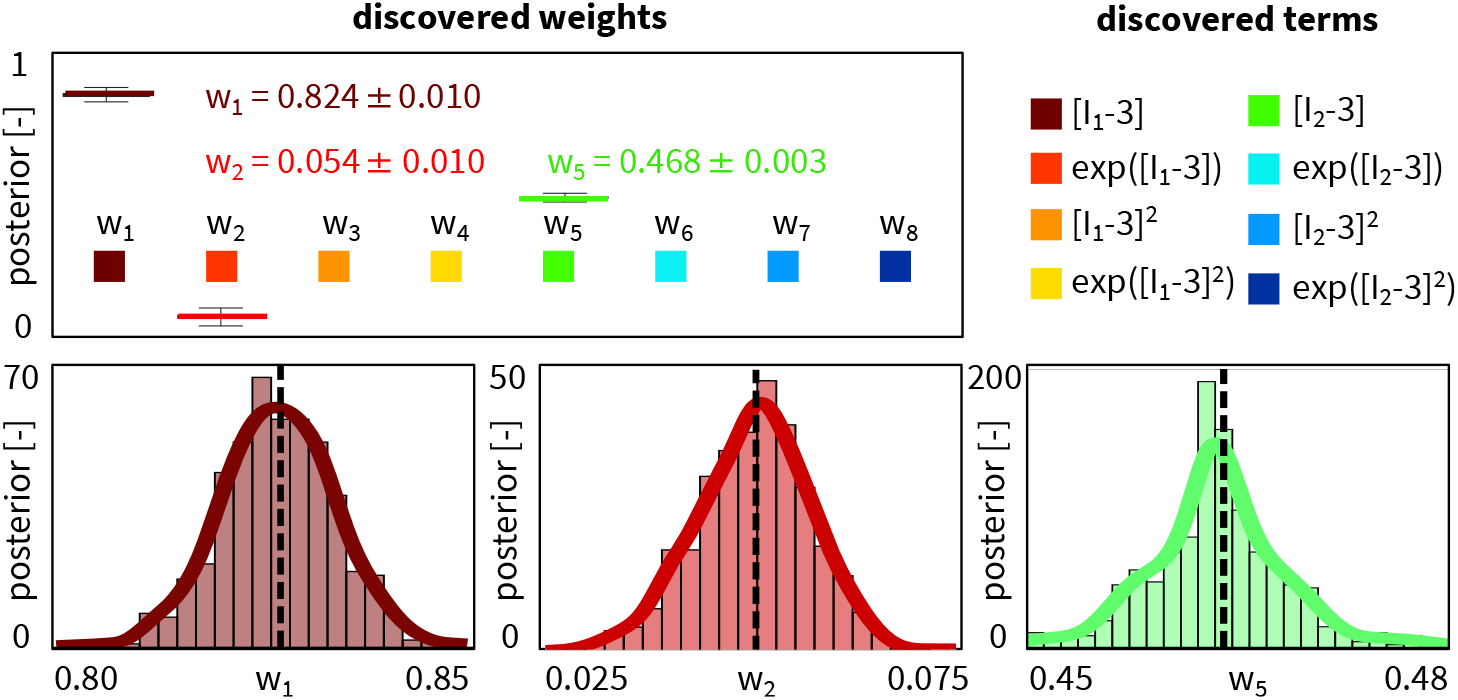
Posterior distributions of discovered isotropic model parameters for training on incomplete data. When using incomplete data for training, the isotropic, perfectly incompressible Bayesian constitutive neural network rediscovers the Gaussian noise perturbed Mooney Rivlin model with an additional term, *ψ* = *w*_1_ [ *I*_1_−3 ]+*w*_2_ [ exp(*w*^∗^[*I*_2_−3])−1 ], +*w*_5_ [ *I*_2_−3 ], with discovered network weights of *w*_1_ = 0.824±0.01, *w*_2_ = 0.054 ± 0.01 and *w*_5_ = 0.468 ± 0.00, while all other network weights correctly train to zero.

## 4. Transversely isotropic Bayesian constitutive neural networks

The *transversely isotropic* Bayesian constitutive neural network takes the deformation gradient ***F*** as input,

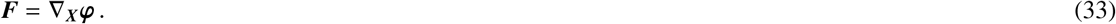

In addition, its kinematics are characterized through a pronounced direction ***n***_0_ with unit length || ***n***_0_ || = 1 in the undeformed configuration, which map onto the pronounced direction ***n*** = ***F*** · ***n***_0_ with stretches || ***n*** || = *λ*_*n*_ in the deformed configuration. We characterize the deformation state through the two isotropic invariants *I*_1_ and *I*_2_, and two anisotropic invariants *I*_4_ and *I*_5_ [39],

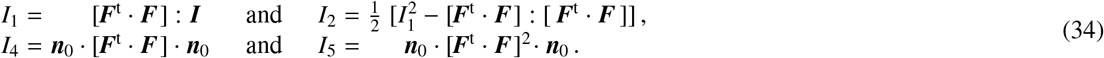

A *perfectly incompressible* material has a constant Jacobian equal to one, *I*_3_ = *J*^2^ = 1. The network discovers *hyperelastic* material models that satisfy the second law of thermodynamics, and their Piola stress ***P*** = ∂*ψ*(***F***)/∂***F*** is the derivative of the free energy *ψ*(***F***) with respect to the deformation gradient ***F*** modified by a pressure term −*p* ***F***^−t^,

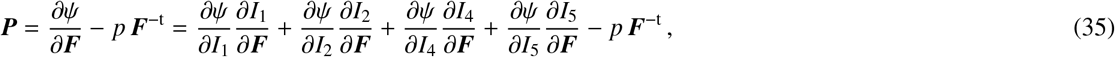

where the hydrostatic pressure, 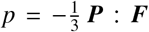, acts as a Lagrange multiplier that we determine from the boundary conditions.

We discover the free-energy function *ψ* using a Bayesian constitutive neural network that takes the deformation gradient ***F*** as input and approximates the free-energy function *ψ*(***F***) as the sum of sixteen probability-weighted terms. Figure 7 illustrates our neural network with two hidden layers and sixteen nodes [23]. The first layer generates powers (○) and (○)^2^ of the network input, the two invariants *I*_1_ and *I*_2_. The second layer applies the identity, (○) and the exponential function (exp(○)) to these powers. The network output is the sum of these sixteen terms, weighted by their probability densities 𝒩(*w*_*µ*_, *w*_*σ*_). The free-energy function of this networks takes the following explicit form,

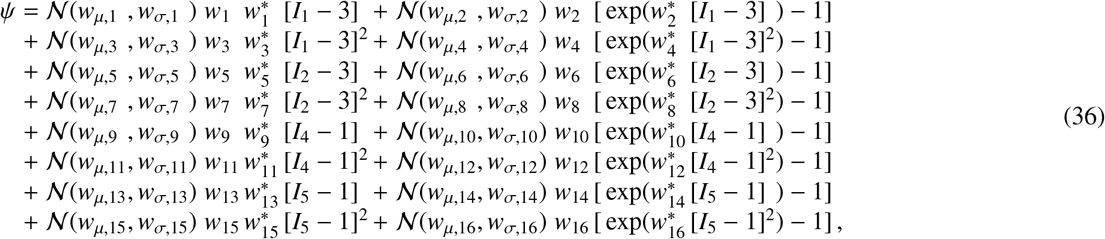

corrected by the pressure term, *ψ* = *ψ* − *p* [*J* − 1]. To complete the definition of the Piola stress in equation (35), we take its derivatives with respect to the two invariants,

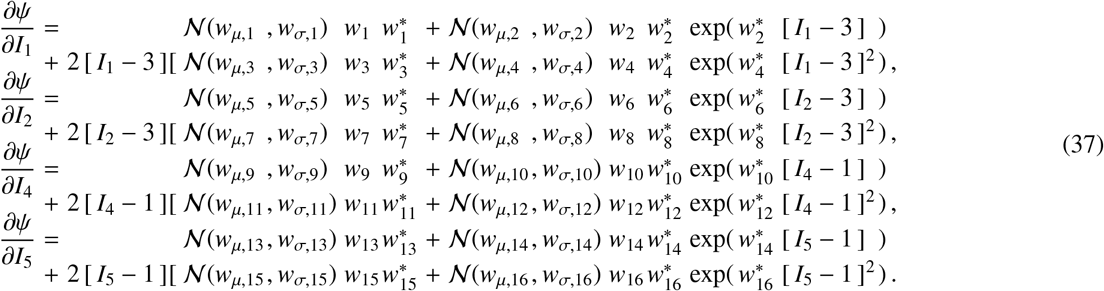

**Figure 7:**
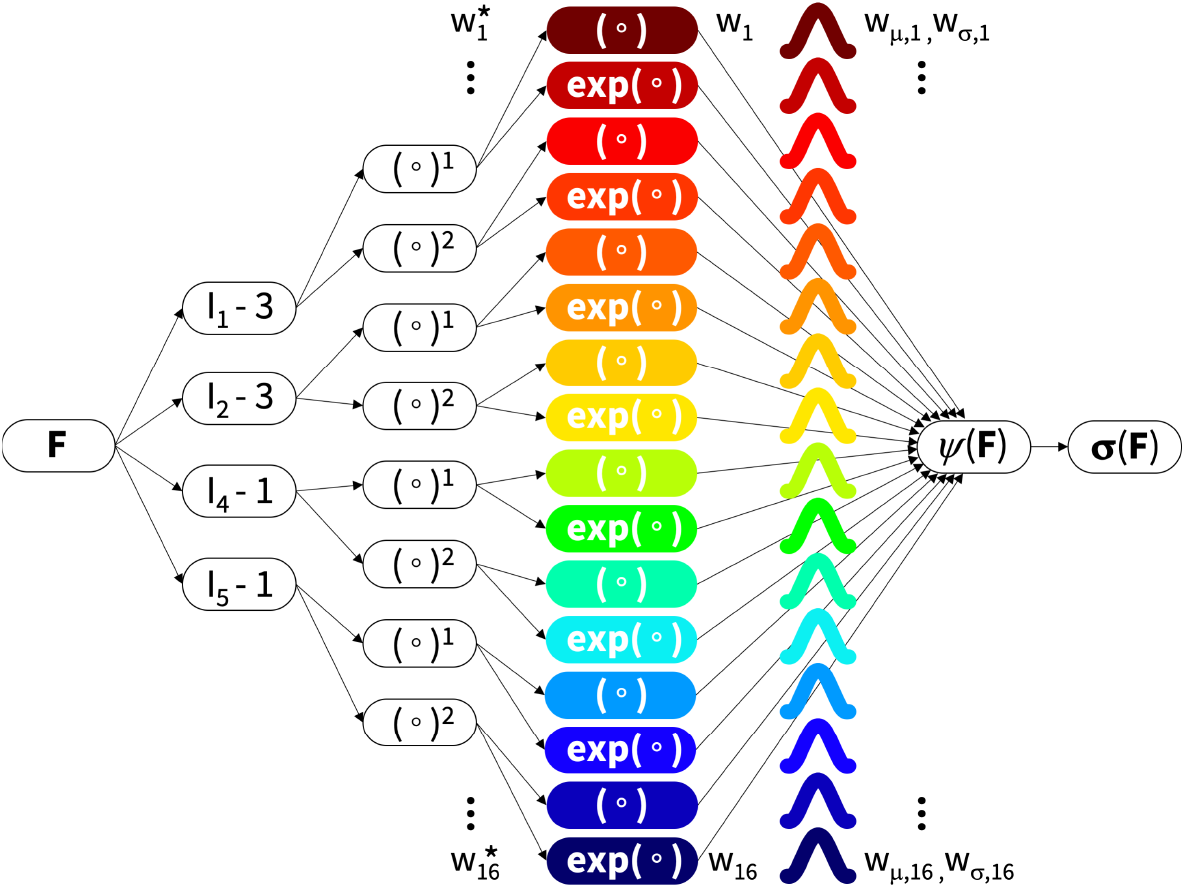
Transversely isotropic Bayesian constitutive neural network. The network has two hidden layers to discover the free-energy function *ψ*(*I*_1_, *I*_2_, *I*_4_, *I*_5_) as a function of the invariants of the deformation gradient ***F*** using sixteen terms. The first layer generates powers (○) and (○)^2^ of the network input, the second layer applies the identity (○) and exponential function (exp(○)) to these powers, and the network output is the sum of these sixteen terms, weighted by their probability densities 𝒩(***w***_*µ*_, ***w***_*σ*_). During training, the network learns the network weights ***w*** and ***w***^∗^ and the variational parameters ***w***_*µ*_ and ***w***_*σ*_.

The network has two sets of network weights, ***w*** and ***w***^∗^, associated the sixteen terms of the free-energy function, and two sets of variational parameters, ***w***_*µ*_ and ***w***_*σ*_, associated with the means and standard deviations of these sixteen terms. We learn these 32 weights by minimizing the three-term loss function *L* = *L*_NLL_ + *L*_KL_ + *L*_1_ in equation (21).

We consider data from *biaxial extension* tests with stretches *λ*_1_ and *λ*_2_ in the 1- and 2-directions, such that 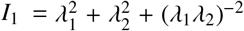 and 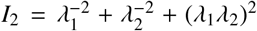and 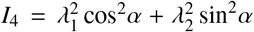and 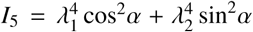. The deformation gradient is **F** = diag { *λ*_1_, *λ*_2_, (*λ*_1_*λ*_2_)^−1^ } and the Piola stress is **P** = diag { *P*_11_, *P*_22_, 0 }. We use the zero-normal-stress condition, *P*_33_ = 0, to determine the pressure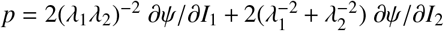 [6]. In what follows, we use data from biaxial extension tests on samples with *two fiber families* with *identical properties* that are mounted symmetrically with respect to the 1- and 2-directions. This implies that we can combine the effects of both fiber families in the fourth and fifth invariants *I*_4_ and *I*_5_ by multiplying the anisotropic stress terms by a factor *n*_f_ = 2. In addition, we assume that these two fiber families do not interact such that the eighth invariant vanishes identically, *I*_8_ = 0 [28]. Equation (35) then provides explicit analytical expressions for the Piola stresses *P*_1_ and *P*_2_ in terms of the stretches *λ*_1_ and *λ*_2_ [27],

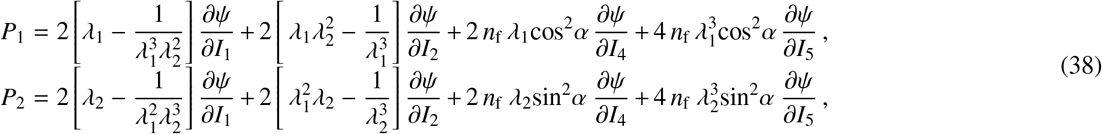

which we translate into the Cauchy stresses *σ*_1_ and *σ*_2_ that are reported in the experiment [31] using *J σ* = ***P*** · ***F***^t^ = ∂*ψ*/∂***F*** · ***F***^t^ − *p* ***I***,

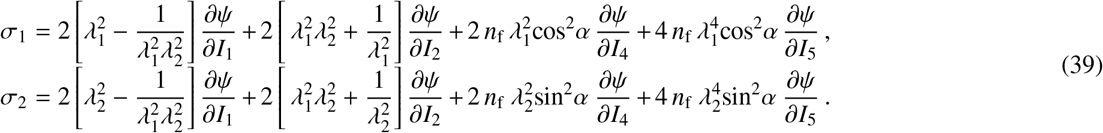

We explore the performance of the transversely isotropic Bayesian neural network from Figure 7 on the basis of real stress-stretch data from human abdominal aortic tissue samples [31, 32]. We consider three data sets of axial and circular stretch stress pairs from the healthy medial layer, the healthy composite aorta, and the aneurysmal composite aorta, in the stretch ranges of 1.00 ≤ *λ*_axl_, *λ*_cir_ ≤ 1.15 and apply Bayesian model discovery: We train the network by minimizing the loss function, draw parameter samples from the learnt variational approximation of the true posterior distribution, derive stresses for each sample, and report the means and standard deviations of the stresses. We compare the discovered models, parameters, and uncertainties.

### Bayesian networks robustly discover model, parameters, and uncertainties for the healthy medial layer

Figure 8, right, shows the equibiaxial extension data and the discovered model for the healthy human medial aorta. The training data consist of axial and circumferential stretch stress pairs from *n* = 7 samples of the medial layer with collagen fiber orientations ***n***_0_ = [cos(*α*), sin(*α*), 0 ]^t^ for *α* = 9.85^○^. Notably, the data points of the healthy medial aorta in Figure 8 display a moderate *inter-sample variation*, a pronounced *stretch stiffening*, and a notable *anisotropy*. For the model discovery, we used *L*_1_ regularization with a regularization parameter *α* = 0.01. Interestingly, out of 2^16^ = 65, 536 possible models, the Bayesian network discovers a familiar model, 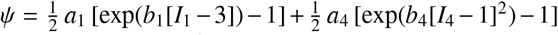, that is dominated by the red exponential linear first invariant Demiray term 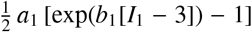, and by the light blue exponential quadratic fourth invariant Holzapfel term [13], 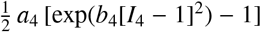 Importantly, the network robustly discovers a *sparse* and *interpretable* model [26], with only two terms–an isotropic term associated with the extracellular matrix and an anisotropic term associated with the collagen fibers–while all other network weights autonomously train to zero. Notably, the discovered model agrees well with a previous analysis of human tissue samples from the healthy medial aorta, which ranked this model within the best-in-class two-term models for the healthy media [24], not only for data from equibiaxial testing, but for data from five different biaxial tests combined [34]. The non-zero weights of the model translate into stiffnesses of *a*_1_ = 1.12 kPa and *a*_4_ = 1.72 kPa and nonlinearity parameters of *b*_1_ = 5.87 and *b*_4_ = 3.96, four meaningful parameters with a clear physical interpretation. Figure 8, left, highlights the discovered model parameters associated with these two terms, *w*_2_ = 1.212 ± 0.11 kPa and *w*_12_ = 1.715 ± 0.09 kPa, along with their posterior distributions and standard deviations. Compared to the previous examples based on synthetic data perturbed by controlled Gaussian noise in Figures 4 and 6, the posterior distributions of the weights in Figure 8, right, display significantly larger standard deviations. This agrees with our expectation of large parameter variations across real biomedical data [31], and with the wide inter-sample spread in the raw data in Figure 8, left. Taken together, our Bayesian constitutive neural network robustly discovers interpretable transversely isotropic models, parameters, and uncertainties from real world data; it not only learns point values, but distributions of parameters with credible intervals, means, and standard deviations that provide valuable information to communicate our confidence in the model, and improve the model if needed.

**Figure 8:**
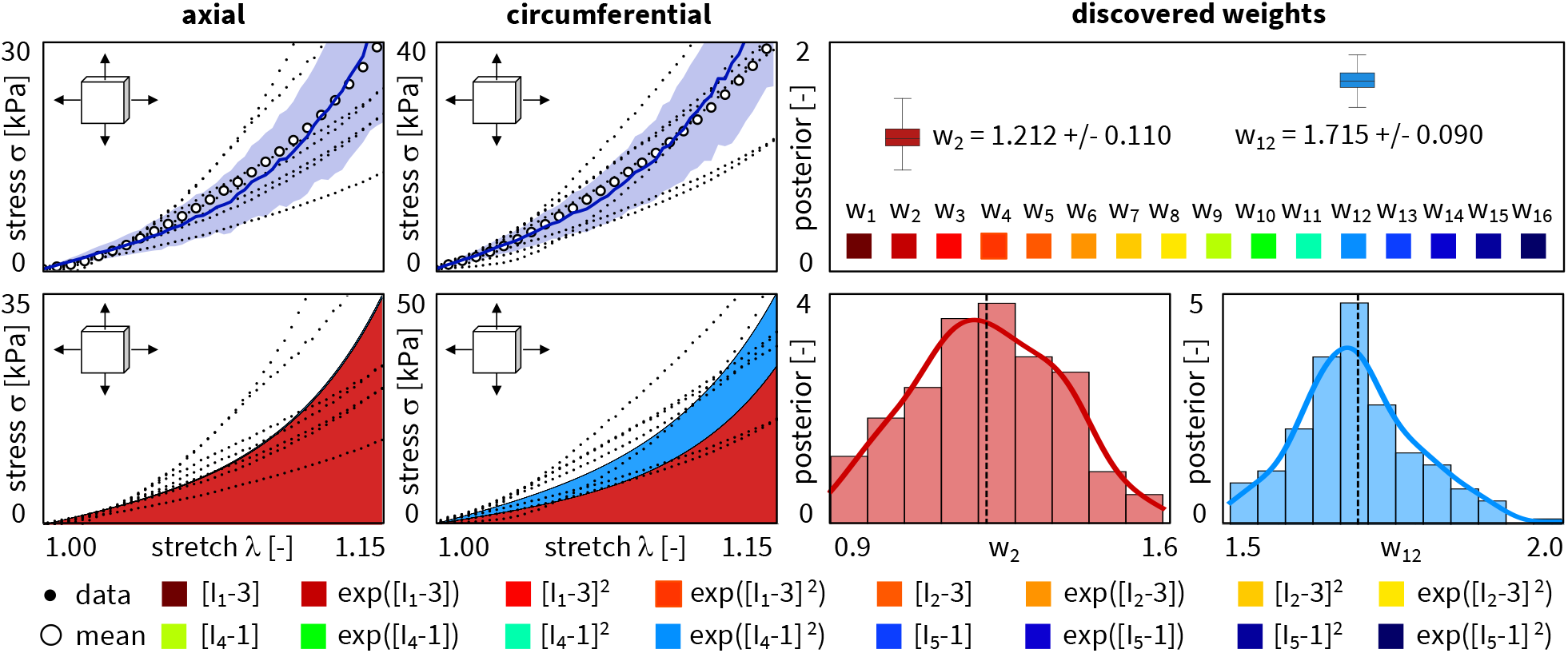
Biaxial extension data and discovered model, parameters, and uncertainties for healthy medial layer. True axial and circum-ferential stresses *σ*_axl_ and *σ*_cir_ as functions of the biaxial stretches *λ*_axl_ and *λ*_cir_ for the transversely isotropic Bayesian constitutive neural network with two hidden layers, sixteen nodes, and sixteen prior distributions from Figure 7. Dots illustrate the axial and circumferential biaxial extension data of *n* = 7 healthy medial aortic samples; solid blue curves and blue-shaded areas indicate the mean predicted stresses ± standard deviations; color-coded areas highlight the sixteen possible contributions to the discovered stress function. The Bayesian network discovers a two-term model, 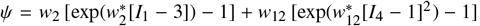, with network weights of *w*_2_ = 1.212 ± 0.11 and *w*_12_ = 1.715 ± 0.09, while all other network weights train to zero.

### Bayesian networks discover a more linear model with more uncertainty for the composite aorta than for the medial layer

Figure 9, left, shows the biaxial extension data and the discovered model for the healthy human composite aorta. The training data consist of axial and circumferential stretch stress pairs from *n* = 6 samples of the composite tissue with collagen fiber orientations ***n***_0_ = [cos(*α*), sin(*α*), 0 ]^t^ for *α* = 29.8^○^. Notably, within the tested stretch regime of 1.00 ≤ *λ*_axl_, *λ*_cir_ ≤ 1.15, and for the tested equibiaxial state, the data points of the healthy composite aorta in Figure 9 display a significant *inter-sample variation*, but *no stretch stiffening*, and only *moderate anisotropy*. As a result, the discovered model, 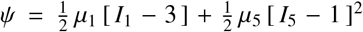, is dominated by the dark red linear first invariant neo Hooke term [42], 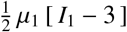 supplemented by the dark blue quadratic fifth invariant term, 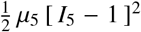. Similar to the previous example, the *L*_1_ regularization promotes a *sparse* and *interpretable* model [26], with only two terms, one isotropic and one anisotropic, while all other network weights autonomously train to zero. This includes the weights of the exponential terms that were prominently featured by the medial layer model [34] to account for the nonlinear stretch stiffening in Figure 8, that is not present in the composite tissue samples in Figure 9. The non-zero weights of the discovered composite model translate into the stiffnesses of *µ*_1_ = 2.748 kPa and *µ*_15_ = 0.362 kPa, two meaningful parameters with a clear physical interpretation. Figure 9, right, highlights the discovered model parameters associated with these two terms, along with their posterior distributions and standard deviations, *w*_1_ = 1.374 ± 0.22 kPa and *w*_15_ = 0.181 ± 0.02 kPa. Again, we observe larger standard deviations than for the synthetic data in Figures 4 and 6, but also larger standard deviations than for the medial layer in Figure 8, right. Confirming this observation, the model uncertainty for the composite aorta associated with the blue-shaded areas in Figure 9, left, is significantly larger than the model uncertainty for the medial layer associated with the blue-shaded areas in Figure 8, left. Since the composite aorta is made up of two mechanically relevant layers [13], the media and the adventitia, both with distinct collagen fiber orientations [31], a mechanistic microstructural model for the composite aorta would need to include at least two anisotropic terms, one for each layer [35]. Instead, we discover a macroscopic phenomenological model that averages the collagen fiber stiffening and anisotropy into a more linear and more isotropic model. Notably, our Bayesian network discovers large aleatoric and epistemic uncertainties that alert us of these unreliable predictions and call for model improvement [33]. Taken together, our Bayesian network autonomously discovers distinct characteristic models for different tissue types: a mechanistic microstructural model with exponential stiffening and notable anisotropy for the medial layer and a phenomenological macrostructural model with a linear response and more isotropy for the composite aorta. Naturally, this composite model contains less microstructural information and generates larger uncertainties.

**Figure 9:**
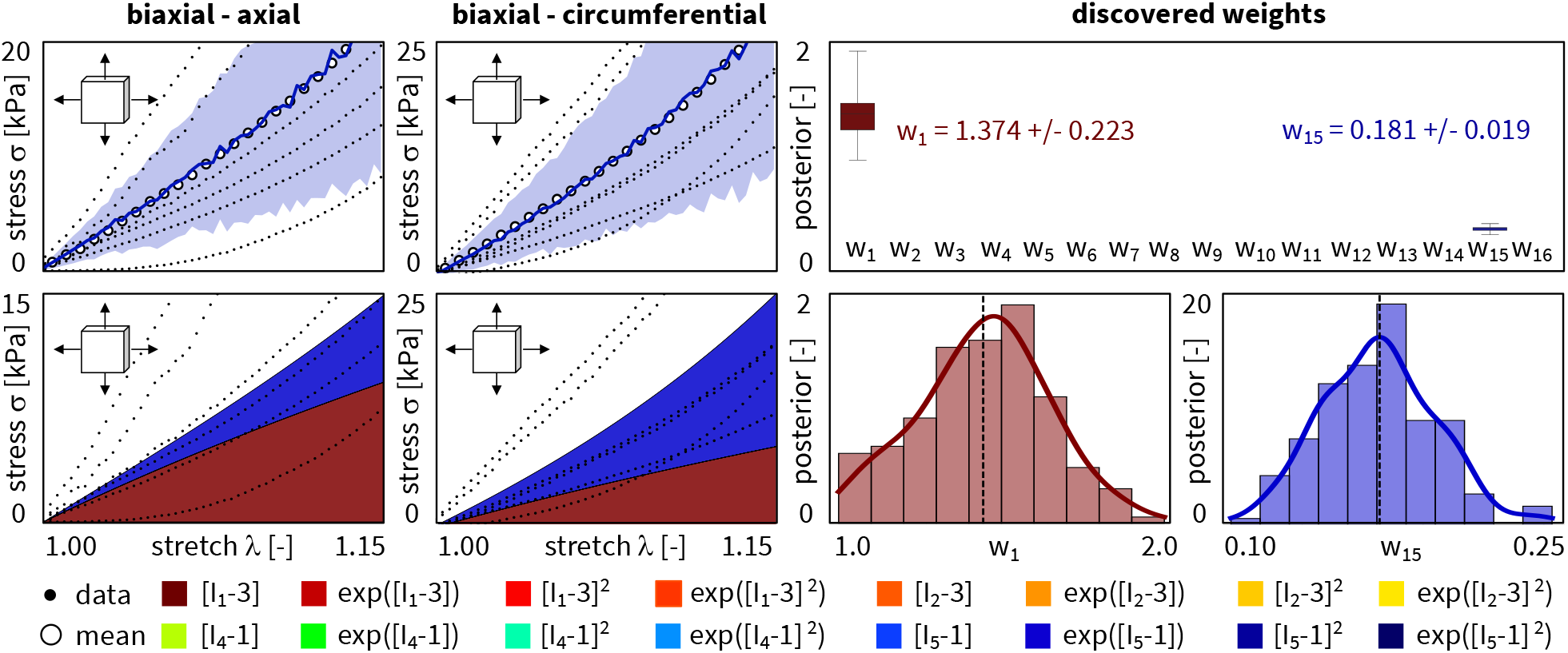
Biaxial extension data and discovered model, parameters, and uncertainties for healthy composite aorta. True axial and circumferential stresses *σ*_axl_ and *σ*_cir_ as functions of the biaxial stretches *λ*_axl_ and *λ*_cir_ for the transversely isotropic Bayesian constitutive neural network with two hidden layers, sixteen nodes, and sixteen prior distributions from Figure 7. Dots illustrate the axial and circumferential biaxial extension data of *n* = 6 healthy composite aortic samples; solid blue curves and blue-shaded areas indicate the mean predicted stresses ± standard deviations; color-coded areas highlight the sixteen possible contributions to the discovered stress function. The Bayesian network discovers a two-term model, *ψ* = *w*_1_ [ *I*_1_ − 3 ] + *w*_15_ [ *I*_5_ − 1 ]^2^, with network weights of *w*_1_ = 1.374 ± 0.22 and *w*_15_ = 0.181 ± 0.02, while all other network weights train to zero.

### Bayesian networks discover a more exponential model with larger uncertainties for the diseased than for the healthy aorta

Figure 10, left, shows the biaxial extension data and the discovered model for the aneurysmal human composite aorta. The training data consist of axial and circumferential stretch stress pairs from *n* = 6 samples with collagen fiber orientations ***n***_0_ = [cos(*α*), sin(*α*), 0 ]^t^ for *α* = 31.5^○^. Similar to the healthy tissue samples in Figure 9, the data points of the diseased tissue samples in Figure 10 display a significant *inter-sample variation*; yet, in contrast to the healthy samples, they also display a pronounced *stretch stiffening*, a notable *anisotropy*. Accordingly, the discovered model, 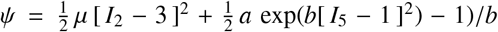, is dominated by the dark blue exponential quadratic fifth invariant Holzapfel-type term [13], 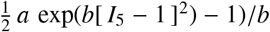, supplemented by a small yellow quadratic second invariant term [18], 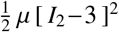 Similar to all previous examples, the *L*_1_ regularized network robustly discovers a *sparse* and *interpretable* model with only two terms, one isotropic and one anisotropic, while all other network weights autonomously train to zero [26]. The non-zero weights translate into the model parameters of *µ* = 0.048 kPa, 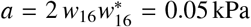, and 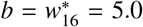. Figure 10, right, highlights the two discovered parameters, their posterior distributions, and their standard deviations, *w*_7_ = 0.024 ± 0.02 kPa and *w*_16_ = 0.042 ± 0.01 kPa. In agreement with our intuition, of all three discovered models, for the healthy medial layer, the healthy composite aorta, and the aneurysmal composite aorta, the model for the aneurysmal composite aorta displays the largest degree of *uncertainty*, as we conclude from the large blue-shaded areas in Figure 10, left. This uncertainty is a natural result of the complex but diffuse microstructure of aneurysmal tissue, dominated by straight and thick struts of collagen [31]. Taken together, our Bayesian constitutive neural networks successfully delineate between healthy and diseased tissues and discover different models–linear for the composite healthy and exponentially stiffening for the composite diseased tissue–both with interpretable parameters, credible intervals, means, and standard deviations.

**Figure 10:**
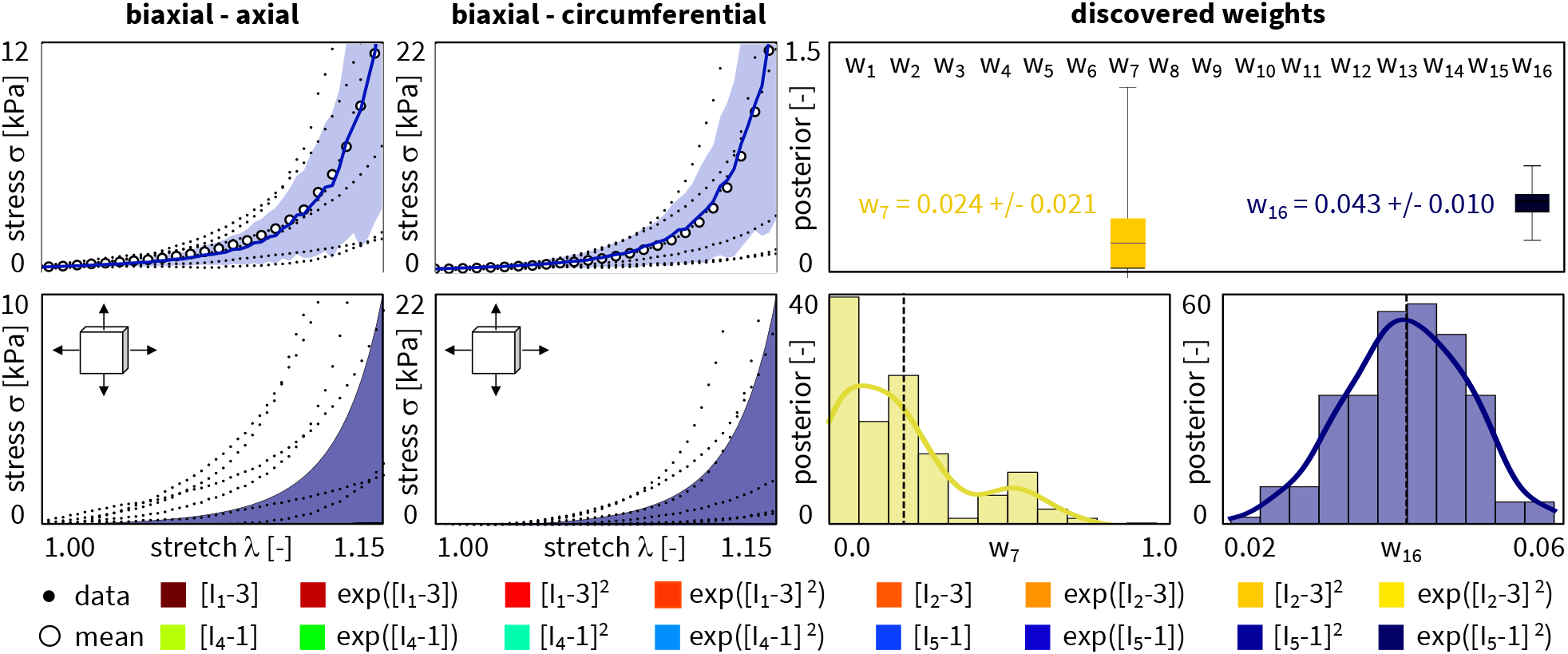
Biaxial extension data and discovered model, parameters, and uncertainties for aneurysmal composite aorta. True axial and circumferential stresses *σ*_axl_ and *σ*_cir_ as functions of the biaxial stretches *λ*_axl_ and *λ*_cir_ for the transversely isotropic, perfectly incompressible Bayesian constitutive neural network with two hidden layers, sixteen nodes, and sixteen prior distributions from Figure 7. Dots illustrate the axial and circumferential biaxial extension data of *n* = 6 aneurysmal aortic samples; solid blue curves and blue-shaded areas indicate the mean predicted stresses ± standard deviations; color-coded areas highlight the sixteen possible contributions to the discovered stress function. The Bayesian network discovers a two-term model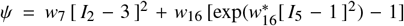, with network weights of *w*_7_ = 0.024 ± 0.02 kPa and *w*_16_ = 0.042 ± 0.01 kPa, while all other network weights train to zero.

## 5. Conclusion

The inability to communicate uncertainty and the risk to produce unreliable predictions are serious deficiencies of classical neural networks. This makes them unsuitable for biomedical applications, where data are sparse and vary significantly from one patient to another. To supplement medical decision making by neural network modeling– especially with a view towards human health–it is absolutely critical to know and understand the uncertainties associated with our model predictions. Here we propose an efficient and robust method, regularized variational Bayesian inference, to train constitutive neural networks and discover the model, parameters, and uncertainties that best explain and predict the unique characteristics of biomedical systems. Importantly, since we focus on model discovery, we only replace the external weight of the network by their probabilistic counterparts, while keeping the internal weights deterministic. This naturally limits the number of additional parameters, and makes the network more robust by design. To demonstrate the potential of this approach, we prototype solutions on synthetic data perturbed by aleatoric noise and on real world data from healthy and diseased human arteries. Our results on synthetic data demonstrate that Bayesian constitutive neural networks can successfully rediscover the initial model, even in the presence of noise, and robustly discover uncertainties, even from incomplete training data. Interestingly, we observe larger uncertainties for equibiaxial tests than for tension, compression, and shear tests. These uncertainties provide valuable guidance for model improvement: If we decided to collect more data, additional equibiaxial tests would most efficiently reduce epistemic uncertainties and improve model robustness. Our results on healthy and diseased human arteries demonstrate that Bayesian constitutive neural networks can successfully discriminate between healthy and diseased tissues, robustly discover interpretable models for both, and efficiently quantify uncertainties in model discovery. Notably, model uncertainty is smallest for the healthy medial layer, moderate for the healthy composite aorta, and largest for the diseased aorta. This observation is in line with an increasing microstructural complexity, from the well-organized healthy medial architecture with pronounced collagen fiber orientations to the distorted aneurysmal tissue with dispersed collagen fragments. Importantly, the failure of the Bayesian prediction presents an opportunity to learn: We could expand our current model library and reanalyze the same data with *new models*, or even collect *new data* and improve model performance in regions with high uncertainties. Especially in risk-sensitive areas like medical diagnosis, it is paramount that we can precisely communicate our confidence in our model predictions. Here we have prototyped this approach for healthy and diseased human arteries. We envision that Bayesian model discovery will generalize naturally to other biomedical systems for which real-world data are rare and inter-personal variations are large. Knowing, understanding, and communicating the uncertainties in automated model discovery is a vital step to improve model prediction, enable personalized simulation, and support informed decision making.

## Data availability

Our source code, data, and examples will be available at https://github.com/LivingMatterLab/CANNs.

## Acknowledgments

This work was supported by the Emmy Noether Grant 533187597 *Computational Soft Material Mechanics Intelligence* to Kevin Linka and by the NSF CMMI Award 2320933 *Automated Model Discovery for Soft Matter* and the ERC Advanced Grant 101141626 *DISCOVER* to Ellen Kuhl.

